# CloneRetriever: retrieval of rare clones from heterogeneous cell populations

**DOI:** 10.1101/762708

**Authors:** David Feldman, FuNien Tsai, Anthony J. Garrity, Ryan O’Rourke, Lisa Brenan, Patricia Ho, Elizabeth Gonzalez, Silvana Konermann, Cory M. Johannessen, Rameen Beroukhim, Pratiti Bandopadhayay, Paul C. Blainey

## Abstract

**Background:** Many biological processes, such as cancer metastasis, organismal development, and development of resistance to cytotoxic therapy, rely on the emergence of rare sub-clones from a larger population. Understanding how the genetic and epigenetic features of diverse clones affect clonal fitness provides insight into molecular mechanisms underlying selective processes. However, identifying causal drivers of clonal fitness remains challenging. Population-level analysis has limited resolution to characterize clones prior to selection, while high-resolution single-cell methods are destructive and challenging to scale across large populations, limiting further functional analysis of relevant clones.

**Results:** Here, we develop CloneRetriever, a methodology for tracking and retrieving rare clones throughout their response to selection. CloneRetriever utilizes a CRISPR sgRNA-barcode library that allows isolation of viable cells from specific clones within the barcoded population using a sequence-specific retrieval reporter. We demonstrated that CloneRetriever can measure clonal fitness of cancer cell models *in vitro* and retrieve targeted clones at abundance as low as 1 in 1,883 in a heterogeneous cell population.

**Conclusions:** CloneRetriever provides a means to track and access specific and rare clones of interest across dynamic changes in population structure to comprehensively explore the basis of these changes.

## Introduction

The response of a heterogeneous population to selection pressure is shaped by the growth dynamics of individual clones within the population. Rare clones can play a decisive role in the outcome of selection. Examples include evasion of anti-retroviral therapy by rare HIV variants [1], expansion of drug-resistant cancer cells under chemotherapy [2], and seeding of metastases by clonal tumor cells [3], [4]. Studying how genetic and epigenetic differences affect the fitness of different clones during selection provides an opportunity to understand both how the selective process operates and how populations are reshaped by selection. In particular, identifying causal drivers of clone fitness could give rich insights into the molecular mechanisms of selection and suggest potential interventions.

Heritable and plastic cellular features can drive selection outcomes. For example, genetic features can change with mutagens, such as DNA-damaging chemotherapies, and epigenetic states can rapidly shift in response to drug exposure [5]or environment [6]. Metastatic clones may alter their epigenetic profiles upon seeding a metastatic site [7], obscuring the preexisting features that enabled them to metastasize. However, existing methods to identify these features tend to rely on comparing populations in bulk before and after selection, which limits their usefulness in detecting pre-existing features that changed during selection. A useful alternative approach would be to identify clones based upon their response to selective pressure, and then isolate representative untreated cells from each clone for genomic and functional characterization.

Genomically integrated DNA barcodes provide a scalable methodology to track rare clones by measuring relative barcode abundance over time [8]. However, relative clone fitness alone cannot elucidate mechanisms of selection. Single-cell technologies can provide genomic profiles of heterogeneous cells within a population. Clone identity can be incorporated into single-cell RNA-seq (scRNA-seq) profiles by capturing transcribed barcodes, linking clonal history and cell fate [9]. However, single-cell genomic profiling is inherently destructive. Both DNA barcoding and single-cell approaches have a limited ability to probe functional differences between clones, whereas retrieval of viable cells from clones would enable a wide range of genomic and functional analysis.

Here, we report CloneRetriever, an experimental system that permits tracking, selection, and recovery of arbitrarily chosen, viable clones from a cell population. CloneRetriever employs a diverse library of single-guide RNAs (sgRNAs). In the absence of Cas9 activity, these serve as inert barcodes for tracking cells. In the presence of Cas9, these sgRNAs direct Cas9 in a clone-specific fashion to activate a reporter. Cas9-dependent reporter expression permits the physical isolation of specific cells within a population while preserving cell viability. This methodology allows for the isolation and comparative analysis of specific clones at any stage of evolution. Isolated cells can then be characterized by downstream functional assays, such as phenotypic characterization, genetic perturbation, or small molecule screens, thus enabling comprehensive analysis of how clonal features affect fitness.

## Results

### Overview of the barcoding and retrieval strategy

To enable tracking and retrieval of clones within a heterogeneous population, we designed a selectable barcode strategy that allows for retrieval of viable cells with clone-specific barcodes. In this system, each clone is tagged with a library of random CRISPR sgRNAs [10]. In the absence of Cas9 expression, the sgRNA-barcodes serve as inert labels that are propagated upon cell division, similar to previously reported clonal barcoding strategies [8],[5]. The relative abundance of each clone can be quantified by deep sequencing of the DNA-integrated sgRNA-barcode. The relative fitness of clones can then be determined by sequencing sgRNA-barcodes over time (e.g., before and after drug selection). By expanding the ancestral barcoded population and splitting the daughter cells into replicate selection assays, clone-specific fitness differences can be estimated (e.g., clones with a drug-dependent fitness advantage) (Figure 1a).

**Figure 1.**
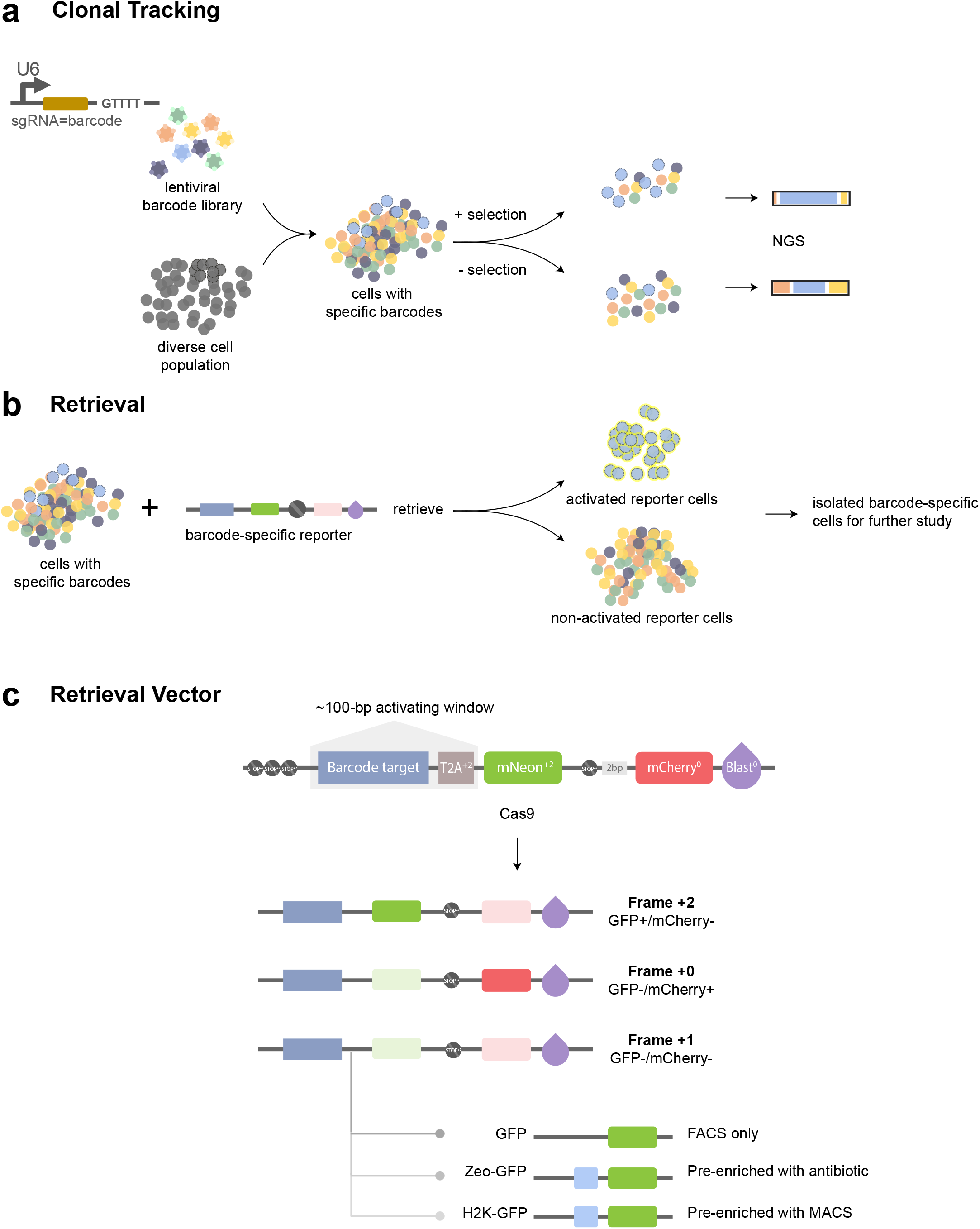
Overview of the strategy for tracking and retrieving the ancestral clones within a heterogeneous population. (a) Tracking clonal response to selection (e.g., ±drug) using a lentiviral sgRNA-barcode library. Clonal fitness profiles can be estimated from barcode enrichment across replicates within each condition. (b) Clones of interest may be retrieved from the ancestral (untreated) population using a retrieval vector containing a targeting region matched to the clone sgRNA-barcode. Nuclease activity at the target region activates a fluorescent marker that can be detected with FACS. (c) Diagram of the frameshift retrieval vector. In cells from the clone of interest, where the sgRNA-barcode and barcode targets are matched, Cas9-mediated cleavage can induce a −1/+2 frameshift, activating reporter expression and inactivating mCherry expression. GFP+/mCherry-cells can be isolated by FACS. Additional reporter genes enable pre-enrichment such as antibiotic selection (e.g., zeocin) or affinity selection (e.g., H2K surface epitope) prior to FACS.

We designed this system so that specific clones can be isolated from a barcoded population using a retrieval vector with a target site matching the sgRNA-barcode of interest (Figure 1b). Introducing Cas9 nuclease leads to double-strand DNA breaks at the target site specifically in the clone that expresses the corresponding sgRNA-barcode. DNA repair generates frameshift mutations at the target site, which may shift the translation frame of one or more downstream reporters [11] (Figure 1c). Activation of the retrieval reporter can result in both gain and loss of reporter expression (e.g., a shift that brings a GFP reporter into frame and an RFP reporter out of frame).

### An sgRNA-barcode library enables tracking clonal subpopulations

We generated two high complexity sgRNA-barcode libraries using fully degenerate oligonucleotide templates of either 20- or 26-nucleotides (nt) (Additional file 1: Fig. S1a). To test the clone tracking capacity of sgRNA-barcodes, we applied the 26-nt barcode library to monitor clonal resistance to the BET-bromodomain inhibitor JQ1, in D458, a MYC-amplified medulloblastoma cell line known to contain pre-existing resistant clones to a chemotherapeutic [5]. We first transduced D458 cells with the 26-nt barcode library at low MOI (< 0.3). We then selected the transduced cells with puromycin and restricted the population size to ensure that a high fraction of barcodes corresponded to unique clones (Figure 2a).

**Figure 2.**
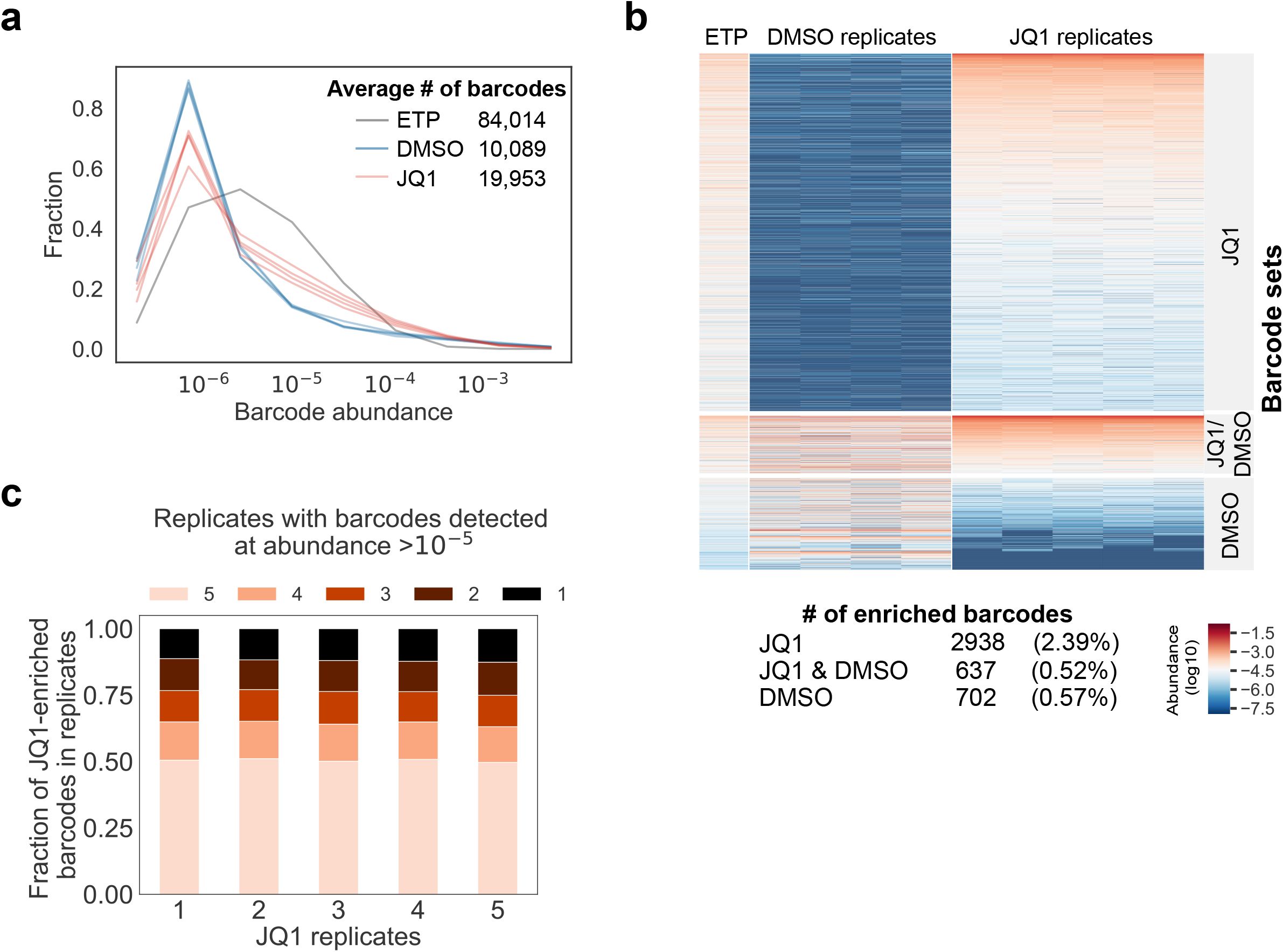
Tracking clonal dynamics in D458 cells using a 26nt sgRNA-barcode library. (a) Relative barcode abundance in D458 cells before treatment (early time point, ETP) and after treatment with 2 uM JQ1 (5 replicates) or DMSO vehicle (5 replicates). (b & c) The sgRNA-barcode library is able to track a heritable phenotype. (b) Comparison of barcode abundance across conditions for barcodes enriched in JQ1, DMSO, or JQ1 and DMSO replicates. Barcode enrichment was defined based on the median rank across replicates (Methods). (c) The majority of JQ1-enriched barcodes were detected across all replicates at an abundance >10^-5^. The raw barcode read counts are provided as a CSV file in Additional file 4: Table S5-barcode_counts.csv and the raw histograms for barcode counts are available in Additional file 6 – barcode_histograms.

We expanded the barcoded D458 population and split it into replicates that were treated with either 2 μM JQ1 or DMSO only (vehicle control). Deep sequencing at the time of the replicate split (early time point, or ETP) detected 84,014 barcodes prior to drug selection (Figure 2a). After 52 days, we harvested cells and quantified barcode abundance in each replicate (Additional file 3: Table S3). An average of 2,938 barcodes were enriched in JQ1-treated replicates, comprising about 2% of the original barcodes. Approximately 50% of the JQ1-selective resistant barcodes were shared by all replicates (Figure 2b and c), suggesting both that these barcodes marked clones with predetermined resistance to JQ1 and that our barcode library enables tracking of clones with such heritable phenotypes within a heterogeneous population. Analysis of barcode enrichments showed no significant biases based on barcode GC content or homology to the genome, suggesting that the sgRNA-barcode library can function similarly to other inert barcoding libraries (Additional file 1: Fig. S1b and c). Although the 26nt-barcoding library enables tracking complex clonal populations, previous studies showed that increasing the length of gRNA sequences above 20nt leads to reduced Cas9 activity [12]. Therefore, we opted to employ a 20-nt sgRNA barcode library that we have shown is also able to track evolution of populations through targeted therapies [13].

### Design of a retrieval vector activated by frameshift mutations

To retrieve viable clones, we designed a frameshift reporter that can be specifically activated by an sgRNA-barcode of interest. This approach relies on the generation of insertion or deletion (indel) mutations by Cas9 nuclease in a target region to shift the translation frame of a reporter cassette, similar to vectors used to monitor gene-editing outcomes [11], [14]. An alternative approach would be to use CRISPR-a (dCas9-transcriptional activator) to activate marker expression in a barcode-dependent fashion [15]. However, we found that a transcriptional activation-based reporter lacked specificity, in part due to a high background level of transcription in a fraction of cells subsequent to lentiviral integration of the reporter (Additional file 1: Fig. S2a and b). Conversely, frameshift reporters have the potential for extremely high specificity due to the low background rate of activating mutations. We opted to deliver the reporter using a lentiviral system, as it can effectively transduce a wide range of cell lines. Lentiviral transduction at low MOI followed by antibiotic selection integrates a single reporter copy into most cells, minimizing the potential for a cell to contain multiple reporters in different frameshift states.

We designed a retrieval vector that gains GFP fluorescence in response to a +2 frameshift mutation that occurs within a narrow targeting window of ∼100 bp. The vector contains two cassettes respectively in the +0 and +2 translation frames: a selection marker (e.g., blasticidin, Blast) and a fluorescent protein (e.g., mCherry) linked by a T2A self-cleaving peptide in the +0 frame, and a second fluorescent protein (e.g., GFP) in the +2 frame. The +0 cassette (mCherry-T2A-Blast) is located downstream of the +2 cassette in order to aid in selecting for integrants with the correct initial frame via antibiotic selection (blasticidin) or fluorescence-activated cell sorting (FACS) (mCherry) (Figure 1c). To minimize the likelihood of background activation, we included triple stop codons in all reading frames immediately upstream of the Kozak translation initiation site. All sequences downstream of the translation initiation site were codon optimized to eliminate start and stop codons that could interfere with reporter performance (Methods).

In order to target a specific barcode, the matching target sequence is cloned into the targeting window between the translation start site and the beginning of the GFP coding sequence. Targeting of Cas9 nuclease by the sgRNA-barcode generates indel mutations in the targeting window. If a +2 indel occurs, the reading frame shifts such that the +0 cassette is out of frame, while the +2 cassette is in frame, giving rise to GFP expression (Figure 1c). In addition to GFP, a variety of alternative selection elements, such as antibiotic resistance or surface affinity markers, can be used to assist in enriching for cells with +2 frameshifts.

### The retrieval vector is specifically activated by target sgRNA-barcodes

We further applied two modifications to improve the activity of our retrieval vector. With the initial version (TMv1), activation with the matching guide produced >1% GFP compared to 0.001% mismatch guide controls (Additional file 1: Fig. S3). To improve sensitivity, we replaced GFP with mNeonGreen and switched the EFS promoter to a stronger EF1a promoter (TMv2). To allow FACS-independent enrichment, we also expanded the +0 selection cassette to include either a Zeocin resistance or H2K surface affinity marker upstream of mNeonGreen (TMv2-Zeo, TMv2-H2K). Compared to TMv1, the TMv2 retrieval vector showed approximately 10-fold increased sensitivity at comparable specificity.

We then systematically evaluated the performance of TMv2 using 5 randomly selected barcodes from our sgRNA-barcode library and matching targets cloned into TMv2 and TMv2-Zeo. We generated HeLa-TetR-Cas9 cell lines expressing each individual sgRNA-barcode, so that specificity and sensitivity could be directly assessed by flow cytometry (Figure 3a). All five barcodes activated mNeonGreen expression from a matching retrieval vector (Figure 3b). In the same experiment, each retrieval vector was tested with mismatched barcode targets to evaluate specificity (Figure 3a). The results showed a low false positive rate ranging from 0 to 2.7⋅10^-5^ for TMv2, and from 0 to 5.5⋅10^-4^ for TMv2-Zeo. The sensitivity for the matched barcodes ranged from 1.8⋅10^-1^ to 2.3⋅10^-2^ for TMv2, and from 0.92⋅10^-1^ to 3.8⋅10^-2^ for TMv2-Zeo, suggesting that the system was capable of high specificity and selectivity.

**Figure 3.**
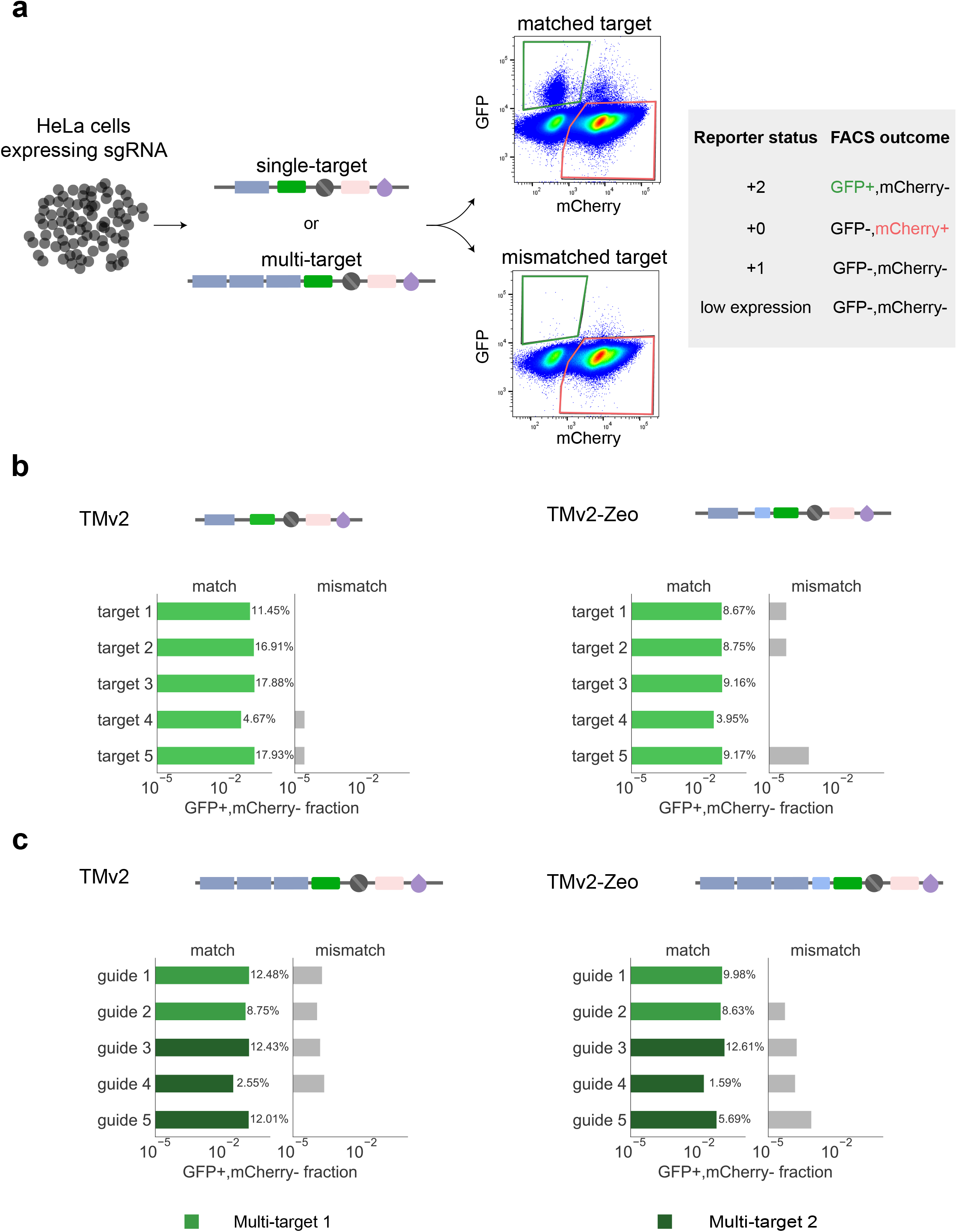
Retrieval vector performance. (a) HeLa cells were transduced with individual sgRNA-barcodes and paired with matched or mismatched barcode targets. Cells with frameshift +2 and 0 are expected to express GFP and mCherry, respectively, whereas cells with a +1 frameshift should express neither. (b) FACS analysis of plots of the TMv2 and TMv2-Zeo retrieval vectors with matched or mismatched barcode targets. (c) Incorporating tandem targets into the retrieval vectors enables multiplexed activation of a single vector by several barcodes. Gating strategy for analysis of the frameshift status of the cells is shown in Additional file 1: Fig. S5. The source data are provided as FCS files in Additional file 8.

In addition to single barcode reporters, multiplexed activation of several barcodes with one reporter can be achieved by expanding the target sequence to contain targets for multiple sgRNA-barcodes (Figure 3a). To demonstrate multiplexing, we designed retrieval vectors to target three independent sgRNA-barcode sequences (Figure 3c). These vectors showed similar sensitivity to those individual sgRNA-barcodes, albeit at 2.6-fold reduced specificity (1.4⋅10^-3^), possibly due to the increased likelihood of background mutations in the expanded target region.

### Identification and viable isolation of rare hygromycin-resistant HeLa cells

We next tested our ability to retrieve drug-resistant and drug-sensitive clones of interest in a well-controlled setting. We engineered hygromycin-resistant HeLa-TetR-Cas9 cells and spiked them into a pool of hygromycin-sensitive HeLa-TetR-Cas9 cells to achieve a final population of cells in which 2% of all cells expressed the hygromycin resistance gene.

We transduced the cells with the 20-nt sgRNA-barcode library at low MOI, and then bottlenecked, expanded, and cryopreserved them in replicate vials. Sequencing of one replicate verified the presence of 441 barcodes ranging in abundance from 1 in 100 to 1 in 100,000 (Figure 4a, Barcoding). To assay for hygromycin resistance, we split the cells and treated them in replicate with either hygromycin or PBS (vehicle) (Figure 4a, Selection). We then nominated candidate hygromycin-resistant barcodes by comparing the abundance of the barcodes in hygromycin–treated cells to the PBS-treated groups (Figure 4a, Deconvolution). We found 15 candidate hygromycin-resistant barcodes with >10-fold enrichment after hygromycin treatment and an ETP frequency of at least 1 in 3,000.

**Figure 4.**
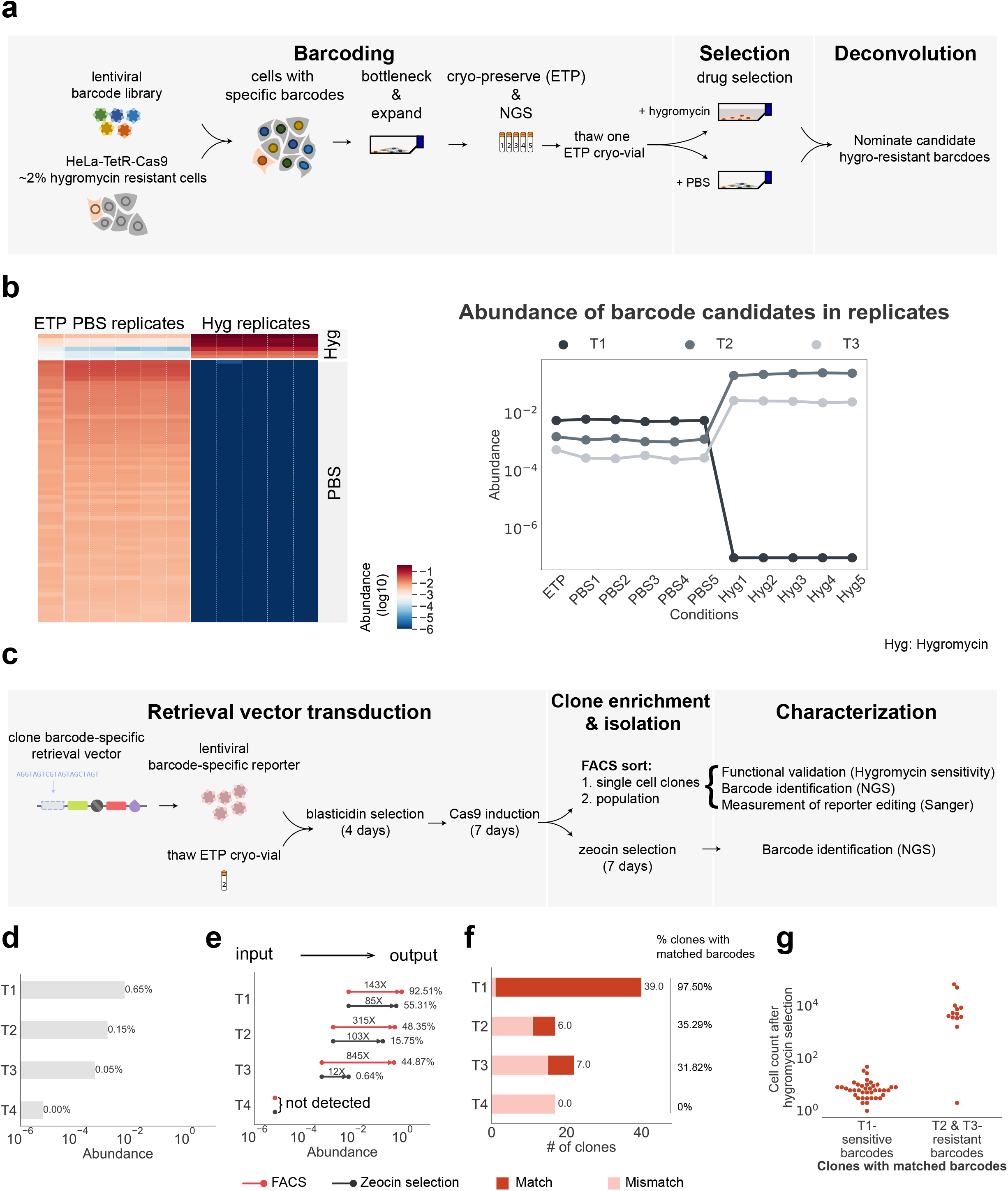
Retrieval of hygromycin-resistant clones from a heterogeneous population of HeLa cells. Workflow to identify resistant clones using an sgRNA-barcode library. (Barcoding) A mixed population of hygromycin-resistant and hygromycin-sensitive HeLa cells was transduced with sgRNA-barcodes. (Selection) The resulting library was bottlenecked to limit barcode complexity, re-expanded, and cryo-preserved to define an early time point (ETP). Cells were then treated with either hygromycin or vehicle control (PBS). Hygromycin-enriched barcodes were determined by NGS. (b) Hygromycin-resistant barcodes were enriched across hygromycin-treated replicates. Barcode abundance for T1 (hygromycin-sensitive barcode candidate), T2 (hygromycin-resistant barcode candidate) and T3 (hygromycin-resistant barcode candidate). The raw barcode read counts are provided as a CSV file in Additional file 4: Table S5-barcode_counts.csv and the raw histograms for barcode counts are available in Additional file 6 – barcode_histograms (c) Workflow to retrieve resistant clones using the frameshift reporter. (Retrieval vector transduction) Hygromycin-sensitive and resistant candidate barcodes were selected for retrieval, and the matching barcode targets were cloned into the retrieval vector. Cells from the ETP were transduced with barcode-specific retrieval vectors and Cas9 expression was induced. (Clone enrichment and isolation) FACS sorting or zeocin selection was used to enrich for barcodes of interest. Single-cell clones were isolated by FACS. (Characterization) Barcode identification and functional validation. The integrated retrieval vector was sequenced to characterize specific and nonspecific mutations leading to reporter activation. (d) The ETP abundance of each targeted barcode. (e) Population-level enrichment of targeted barcodes using selection by FACS (TMv2) or Zeocin selection (TMv2-Zeo). (f) Fraction of single-cell clones with the targeted barcode. (g) The hygromycin sensitivity of single-cell clones isolated by FACS corresponded to the sensitivity predicted by clonal tracking.

We carried out retrieval for 4 clonal barcodes: one hygromycin-sensitive barcode candidate (T1) and 3 hygromycin-resistant barcode candidates (T2, T3 and T4) that were represented in the population at frequencies ranging from 1 in 652 (T2) to 1 in 140,000 (T4) (Figure 4b and d) (Additional file 1: Table S1 and Additional file 4: Table S2). We also analyzed 2 types of control populations: cells transduced with retrieval vectors targeting barcodes not present in the library, and cells without doxycycline induction of Cas9.

For each sgRNA-barcode, we cloned a matching targeting sequence into the retrieval vectors TMv2 and TMv2-Zeo. To retrieve cells representative of the initial, unselected population, we thawed and expanded barcoded cells preserved at the ETP. Barcoded cells were transduced with either TMv2 or TMv2-Zeo and selected with blasticidin for 4 days. Blasticidin was then removed and Cas9 expression was induced with doxycycline for 7 days (Figure 4c, Retrieval vector transduction).

This process greatly enriched clones T1-T3. FACS purification followed by expansion and sequencing indicated up to 845-fold enrichment of these clones relative to the ETP fraction, to a minimum purity of 44.87% (T3) and a maximum purity of 92.51% (T1) (Figure 4d and e, FACS). In addition to the FACS-based enrichment, we also carried out selection using TMv2-Zeo, and detected 12 - 85-fold enrichment (Figure 4e, Zeocin selection). Clone T4 was not present in the enriched population, suggesting the sensitivity of the retrieval vectors was insufficient to recover viable clones present at frequencies in the population that are smaller than 1 in 140,000 (Figure 4e).

### Validation of retrieved clones and analysis of sensitivity-limiting background events

In order to confirm that the barcoding and retrieval protocols led to the recovery of clones exhibiting the hygromycin-resistant phenotype, we sorted individual cells transduced with TMv2 into multi-well plates and expanded them as clonal populations (Figure 4c, Clone enrichment & isolation). We analyzed a total of 132 single-cell clones by deep sequencing of their sgRNA-barcodes. We detected 2 populations: clones with exact matches to the targeted barcodes (52/132 clones), and mismatched clones (barcode edit distance >9; 80/132 clones) (Figure 4f and Additional file 1: Fig. S4a, red square). The hygromycin sensitivity of the single-cell clones reflected their barcodes, with the exception of one single-cell clone with a candidate hygromycin resistance barcode (T3) that was sensitive to hygromycin (Figure 4g and Additional file 1: Fig. S4a, blue square).

To investigate activation of the retrieval vector, we performed Sanger sequencing of 75 clones over a 2-kb region encompassing the translation start site, barcode-specific targeting region, and mNeonGreen coding sequence. As expected, clones with the correct sgRNA-barcode contained +2 frameshift mutations in the targeting region, with a distribution of indel sizes consistent with repair by non-homologous end-joining following Cas9 cleavage (Additional file 1: Fig. S4c) [16]. In contrast, 21/43 false positive clones exhibited a ∼80-nt stereotyped deletion immediately upstream of the mNeonGreen coding sequence (Additional file 1: Fig. S4b and c). Deletions due to lentiviral intra-molecular recombination between homologous regions are well-characterized [17].

However, our codon-optimized retrieval vector lacks substantial homology near the deleted region (no repeated kmers with length >7), suggesting an alternative mechanism. The false positive events observed were largely due to the stereotyped deletion, as we found that sorting error likely did not contribute to these false positive events, as re-analysis of expanded clones showed that all clones contained GFP+/mCherry-cells (Additional file 1: Fig. S4a, green square).

Together, these results indicate that CloneRetriever is capable of tracking hygromycin-resistant phenotypes under treatment, and enriching rare clones up to 800-fold.

## Discussion

We engineered a molecular tool that couples an sgRNA-barcode library for tracking clones with a Cas9-based frameshift reporter to isolate viable cells representing target clones from the population. A challenge to studying the mechanisms underlying clonal evolution has been that bulk methods have limited resolution to observe characteristics of rare clones, while single-cell methods have not been able to target clones with known evolutionary paths. We showed that our system can accurately track clonal fitness under drug selection and allows efficient retrieval of a targeted set of clones at frequencies as low as 1 in 1,883. We demonstrate that a CRISPR sgRNA-barcode approach is able to scale to high complexity libraries capable of barcoding >10^5^ clones, while a frameshift retrieval reporter activated by barcode-specific Cas9-mediated mutations enables fluorescence-based retrieval of clones. The sgRNA-barcode design is especially conducive to multiplexing for simultaneous retrieval of a handful of clones at a time (as shown in Figure 3c), because it allows straightforward expansion of the activating window to accommodate multiple sgRNA-barcodes.

Isolating clonally barcoded cells from an untreated, ancestral population enables direct testing of mechanisms underlying differential clone fitness. Unlike bulk methods that rely on strong positive selection to enrich for cells of interest, our method allows retrieval from clones with any fitness profile, such as slow-growing, persistent, or negatively selected clones. The ability to expand pure populations of target clones enables the use of a broad range of functional and molecular profiling assays. For example, access to pure populations enables high-input assays to determine how epigenetic alterations, such as changes in DNA methylation and chromatin state, affect fitness differences between genetically similar clones. Deep characterization of purified resistant clones can be useful in identifying resistant drivers, and through perturbational approaches, the association between these putative drivers and phenotype can be defined. The sgRNA-barcodes can also be readily adapted to existing high-throughput single-cell readouts developed for CRISPR screens, such as single-cell gene expression [18], [19] and optical screening [20].

An important caveat for this and other lentiviral-based DNA barcoding strategies is the possibility of unintended side effects from semi-random lentiviral integration on barcoded clones. Lentiviral integration can either disrupt or increase gene activity, leading to clone-specific effects. Our approach, in which clones of interest are isolated, simplifies sequencing of the DNA barcode insertion site, which can help rule out integration-driven effects. While introducing the retrieval vector requires an additional lentiviral integration event, multiple independent sub-clones can be retrieved per clone of interest, serving as biological replicates for the retrieval process. A second potential confounding factor is sgRNA-barcode sequence-specific effects on clone fitness in the absence of Cas9. However, we found no significant correlation between enrichment in a clone tracking experiment and sgRNA-barcode sequence content or sgRNA homology to the genome.

Background activation of the frameshift retrieval reporter may hinder applications where the clones of interest exist at low frequency. In principle, a reporter activated by an indel mutation in a 100-nt activating window could have a background rate as low as ∼1 in 1 billion per cell division, as the rate of naturally occurring indel mutations in human cells is estimated to be ∼1 in 10^11^ indel/bp/cell division per generation [21]-[23]. We identified stereotyped deletions in the T2A linker region as the primary source of false positive activations; optimizing this sequence could significantly suppress background. Alternatively, negative selection against the GFP-containing frame could be applied prior to editing to remove cells with a premature frameshift. To improve sensitivity to both +1 and +2 frameshifts, a second reporter cassette could be added in the +1 frame. Selecting sgRNA-barcodes based on their predicted indel distribution could further increase activation efficiency [24].

## Conclusions

Clone tracking and retrieval enable deep, mechanistic studies in a wide range of selection scenarios. For example, tracking cells during reprogramming or differentiation protocols would enable isolation and epigenetic characterization of ancestral clones that are predisposed to successful outcomes [9], [25]. Similarly, retrieving untreated cells from clones surviving mutagenic chemotherapy, such as alkylating agents, could address outstanding questions about whether resistance is pre-existing or acquired [26]. Clones can also be targeted based on fitness profiles derived from multiple parallel selection conditions. Altogether, live clone retrieval capability enables barcoding experiments to advance from observing clone frequency statistics toward experimentally-driven mechanistic studies by providing access to key samples supporting a wide range of genomic and functional assays.

## Methods

### Library construction

Degenerate oligos for sgRNA-barcode library construction were synthesized by IDT and cloned into lentiGuide-Puro [27] by Gibson assembly as previously reported [28]. Approximately 300 ug of Gibson product was transformed into 25 uL of Endura electrocompetent cells (Lucigen). After a 1 hour recovery period, 0.1% of transformed bacteria were plated in a 10-fold dilution series on ampicillin plates to determine the number of successful transformants. The remainder of the transformed bacteria were cultured in 50 mL of LB with 50 ug/mL ampicillin for 16 hours at 30° C. Plasmid libraries were extracted using Plasmid MidiPlus kit (Qiagen) and sequenced to a depth of 95 million reads on Illumina Nextseq, corresponding to 13X coverage of 3.9 million barcodes. Lentivirus was prepared as previously reported [28] by transfecting a total of 10 million HEK 293FT cells. The library virus was determined by transduction and puromycin selection in HeLa-Tet-Cas9 cells to contain 600 million infective particles, corresponding to a 153X coverage of barcodes.

### Barcoding of HeLa and D458 cell lines

HeLa-Tet-Cas9 cells were cultured in DMEM medium supplemented with 10% tetracycline-screened FBS (Hyclone) and 1% penicillin-streptomycin. sgRNA-barcodes were transduced as previously described [28] and selected with 1 ug/mL puromycin for 3 days. The lentiviral multiplicity of infection (MOI) was determined to be between 0.05 and 0.3 for all libraries, so that a majority of cells carry a single integrated sgRNA-barcode. Barcoded cell lines were expanded to a total of 1.0⋅10^7^ cells and cryopreserved in aliquots of 1.0⋅10^6^ cells for subsequent drug selection and retrieval. D458 medulloblastoma cells were cultured in DMEM/F12 media supplemented with 10% FCS and 1% GPS (glutamate, pen-strep). Four million cells were transduced with the sgRNA barcode library (10 wells of 3.0⋅10^6^ cells with 50ul of virus) by spin infection (1,000g, 120 minutes, 30° C). Selection with 1 ug/mL puromycin was initiated 48 hours post-transduction and maintained for a total of 3 days.

### Drug resistance experiments: D458 and JQ1

Barcoded D458 medulloblastoma cells (fingerprint verified) were treated with DMSO or JQ1 at a concentration of 2uM in multiple replicate plates (5 x DMSO and 5 x JQ1). Four million barcoded D458 cells were plated in each replicate plate in presence of DMSO or JQ1. Barcoded D458 cells were also frozen in 10% DMSO/FCS for future retrieval. In addition, cells were collected for DNA-extraction to determine barcode representation at the early-time point (ETP). Cells were retreated with compound every 3-4 days. Cells were counted and passaged every 3-4 days, maintaining a minimum representation of 4 million cells. Cells were cultured in DMSO or JQ1 for a total of 52 days prior to harvesting for DNA extraction for barcode sequencing and deconvolution.

### Drug resistance experiments: HeLa and hygromycin

HeLa-TetR-Cas9 cells were infected with a lentiviral ORF construct (pLX_TRC317_PGK-Hygro) containing a hygromycin resistance cassette. After selection with 300 ug/ml hygromycin for 1 week, HeLa-LacZ cells were spiked into uninfected cells at a ratio of 1:50. Cells were then infected with the CloneRetriever library at MOI <0.3. Following selection with puromycin, we plated a fixed number of cells (to achieve a ‘bottleneck’ of the number of barcoded cells) and expanded the population. Cells were frozen in liquid nitrogen (early time point, ETP) in replicates of 1⋅10^7^ cells. One replicate was thawed for barcoding experiments (1 x ETP, 5 x DMSO and 5 x hygromycin at 300 ug/ml). Replicate cells were cultured in DMSO or hygromycin for 16 days, after which DNA was extracted from both the ETP control and DMSO/hygromycin treated replicates for barcode sequencing and deconvolution. At each passage, we ensured the number of cells plated was at least 10-fold the library complexity in order to maintain representation.

### Library deconvolution

Genomic DNA was extracted and prepared for deep sequencing as reported [28]. Libraries were sequenced to a minimum depth of 18 million reads, corresponding to a barcode coverage of >80X. Counts of sgRNA-barcodes were obtained by filtering for reads containing exact matches to the flanking sequences, and matches with <3 reads were discarded.

### Clonal fitness measurements

Relative clone abundances were calculated from normalized read counts and clones were ranked by abundance within each replicate. For the D458 clonal tracking experiment, JQ1- and DMSO-enriched barcodes were defined as those with a median rank above 4,000 in JQ1 replicates and above 2,000 in DMSO replicates. NGS data analysis were run with Python 2.7 with its libraries numpy 1.13.1, matplotlib 2.1.2, seaborn 0.9.9, and jupyter 4.3.0.

### Retrieval reporter construct

The mNeon, T2A, Zeocin, H2K, and Blasticidin coding sequences were codon optimized with silent nucleotide substitutions to remove out-of-frame start and stop codons. Oligos containing targeting barcode sequences and PAM (NGG) matching barcodes of interest were synthesized (IDT) and cloned into frameshift reporter plasmids by golden gate assembly. All targeting barcode sequences were filtered to have <70% GC content, no more than 4 consecutive repeated bases and no stop codons. Lentivirus was prepared as previously described [28] and transduced into barcoded HeLa-Tet-Cas9 cells at an MOI of <0.3. After 4 days of selection with 10 ug/mL blasticidin, 1 ug/mL doxycycline was added to induce Cas9 expression. Cells were harvested for deep sequencing as previously reported [28].

### FACS sample preparation and analysis

HeLa cells were carefully washed with PBS and trypsinized with TrypLE Express (Gibco) for 5 minutes. DMEM media contained 10% FBS and 1% Penicillin/Streptomycin was used to neutralize trypsin prior to FACS analysis. Fluorescent protein expression was measured on a Cytoflex flow cytometer. FlowJo V10 was used for analysis. Populations were sorted with high-purity mode on a SONY-SH800 FACS machine, and expanded for 2 weeks before deep sequencing. All analyzed populations were first gated on FSC-A/FSC-H and FSC-A/SSC-A to identify singlets and cells respectively (Additional file 1: Fig. S5).

### Characterization of clones

FACS-sorted clones were trypsinized in plate 3 days after sorting and further expanded for ∼7 days. GFP or mCherry expression for each clone was validated on a Cytoflex flow cytometer. To determine hygromycin sensitivity, the clones were treated with or without 300 ug/ml hygromycin and the media was replenished with fresh hygromycin every 3 days for 7 days. Cell number was measured with a Cytoflex flow cytometer. For Sanger analysis, a 2-kb region of the lentiviral transgene was PCR-amplified from the EF1a promoter (forward strand, primer pTM_negative_fwd) and from the Blast gene (reverse strand, primer pTM_negative_rev) and sequenced with sanger sequencing primer (pTM_sanger_primer) (Additional file 1: Table S2).

### Analysis of frameshift status and indel calculation

For each clone, we used the corresponding unedited retrieval vector as a reference sequence for alignment of Sanger sequencing traces. We determined the location of insertion/deletion/substitution mutations by manual inspection and summarized the mutation as follows. The mutation length (d) was calculated as the difference between the length of the Sanger sequenced vector and the reference sequence, restricted to a window defined by high Sanger quality. The frameshift status was defined as (d) modulo 3. To identify the indel location and length, we focused on the region between the translational start site and the mNeonGreen coding sequence. We then identified the first (reporter.prefix) and last (reporter.suffix) bases of the prefix and suffix sequences of the edited retrieval vector and the first (reference.prefix) and last (reference.suffix) bases of the prefix and suffix sequences of the corresponding region of the reference locus. We then defined ‘query gap’ and ‘reference gap’ as the difference between the prefix and suffix bases of the edited retrieval vector and the reference locus, respectively. (query gap = reporter.suffix - reporter.prefix; reference gap = reference.suffix - reference.prefix). The overall indel outcome was considered an insertion if the query gap exceeded the reference gap; otherwise, it was considered a deletion.

### Cell line authentication

HeLa-TetR-Cas9 cells were a gift from Iain Cheeseman (MIT, Whitehead Institute). D458 cell-lines were a gift from Dr. Bigner (Duke University). To ensure the authenticity of cell lines, we performed Fluidigm SNP-based fingerprinting of each model cell line prior to screening. Cells were routinely tested to exclude the presence of mycoplasma.

## Declaration

### Availability of data and materials

The barcode read counts table for Figure 2 and Figure 4 are available in Additional file 4 – Table S5-barcode_counts.csv. Python scripts used for NGS analysis are available in Additional file 5 – Barcode count dataframe.ipynb and Additional file 7 – paella Python module. The raw histograms for barcode counts are available in Additional file 6 – barcode_histograms. FCS files containing flow cytometry data supporting the conclusions of Figure 3 are available in Additional file 8 – Figure 3. Sanger sequencing datasets supporting the conclusions of Fig S4 are available in Additional file 9 – Sanger sequencing files.

### Competing interests

R.B. and P.B. receive grant funding from the Novartis Institute of Biomedical Research for an unrelated project. C.M.J. is currently a full-time employee and stockholder of Novartis Institutes of BioMedical Research, Inc. The Broad Institute, Dana-Farber Cancer Institute, and MIT may seek to commercialize aspects of this work, and related applications for intellectual property have been filed.

### Funding

This work was supported by SPARC funding from the Broad Institute (R.B. and C.M.J.), a grant from the Bridge Project of the Koch Institute for Integrative Cancer Research at MIT and the Dana-Farber/Harvard Cancer Center (R.B. and P.C.B.), the St Baldricks Foundation (P.B.), the Pediatric Brain Tumor Foundation (P.B. and R.B.), Alex’s Lemonade Stand Foundation (R.B.), an NIH K99 award CA201592-02 (P.B.), NIH 1U54CA224068-01 (C.M.J.), NIH R01 awards CA188228 (R.B.), CA219943 (R.B. and C.M.J.), and HG009283 (P.C.B.), the Jared Branfman Sunflowers for Life Fund for Pediatric Brain and Spinal Cancer Research (P.B. and R.B.). P.C.B. is supported by a Career Award at the Scientific Interface from the Burroughs Wellcome Fund.

### Author contributions

D.F. and A.J.G. developed the frameshift retrieval vectors. D.F., F.T., A.J.G performed clone retrieval and characterization. D.F., F.T., A.J.G., R.O.R., L.B., P.H., E.G., and P.B. performed experiments. D.F. and F.T. analyzed data. P.C.B., C.M.J., R.B., and P.B. supervised the research. D.F., F.T., P.C.B., C.M.J., R.B., and P.B. wrote the manuscript with contributions from all authors.

## Acknowledgements

We thank Emily Botelho and members of Blainey lab for feedback and discussions and acknowledge the Broad Flow Cytometry Facility for experimental assistance.

## Additional files

Additional file 1: Fig S1-S5 and Table S1 and S2 (DOCX, 1.4 MB),

Additional file 2: Table S3 (XLSX, 8.8 MB),

Additional file 3: Table S4 (XLSX, 299.8 KB)

Additional file 4: Table S5-barcode_counts (CSV, 7.6 MB),

Additional file 5: Barcode count dataframe (IPYNB, 5.3 KB).

Additional file 6: barcode_histograms (HIST.zip, 5.9 MB)

Additional file 7: Python paella module (PY, 131 KB)

Additional file 8: Figure 3 (FCS, 1.82 GB)

Additional file 9: Sanger sequencing files (SEQ, 393 KB)

## Supplementary Figures

**Fig. S1.**
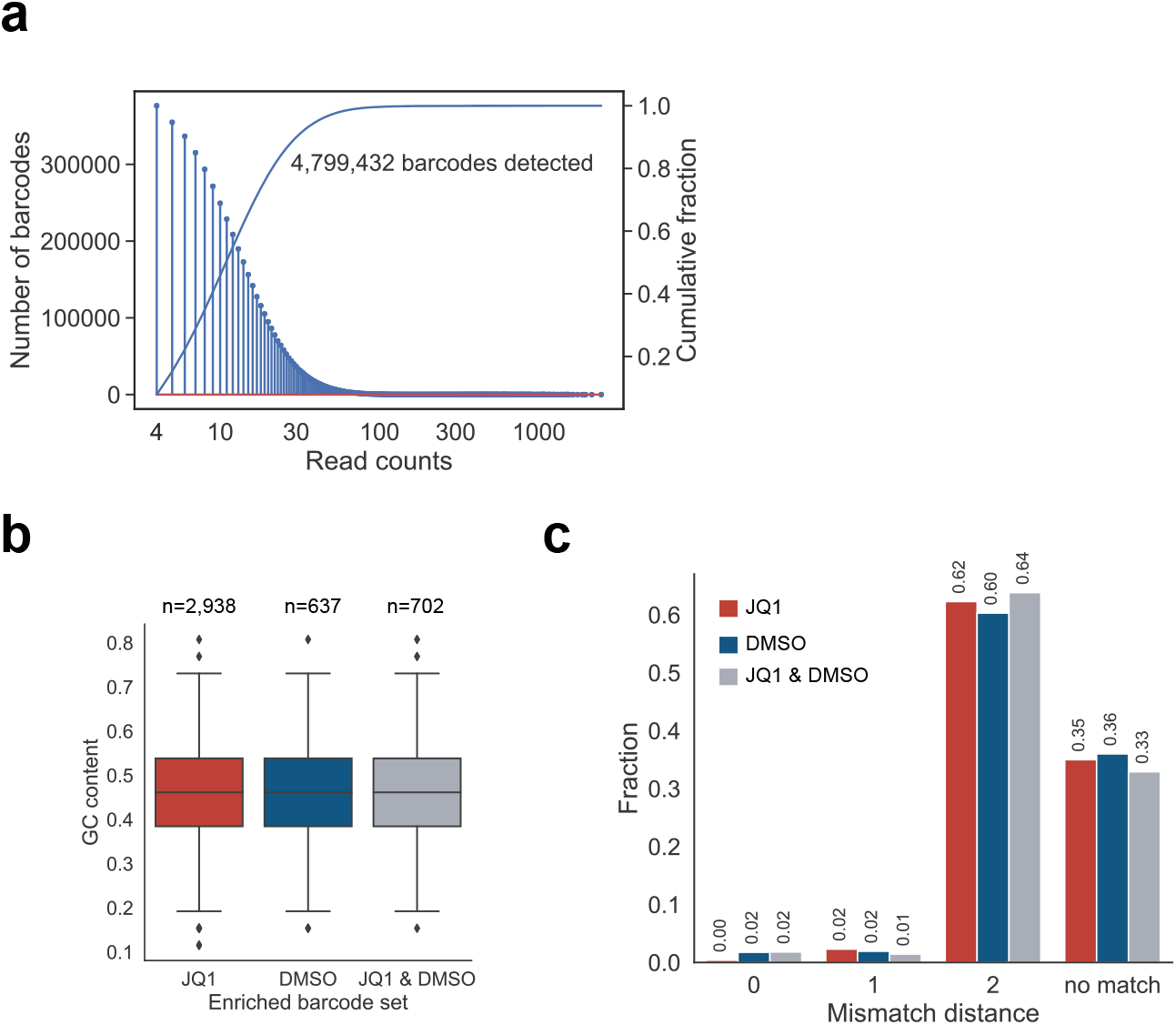
sgRNA-barcode library representation, GC bias and human genome off-targets. (a) Deep sequencing of the 26-nt sgRNA-barcode library plasmid pool. (b) GC content in the 26-nt sgRNA-barcode sequence. There is no significant difference between the barcode sets. (p = 0.07, one-way ANOVA) (c) Distance of sgRNA-barcodes to human genome predicted by an off-target sgRNA algorithm [1]. The vast majority of sgRNA-barcodes have mismatch distance ≥ 2 homology to human genome.

**Fig. S2.**
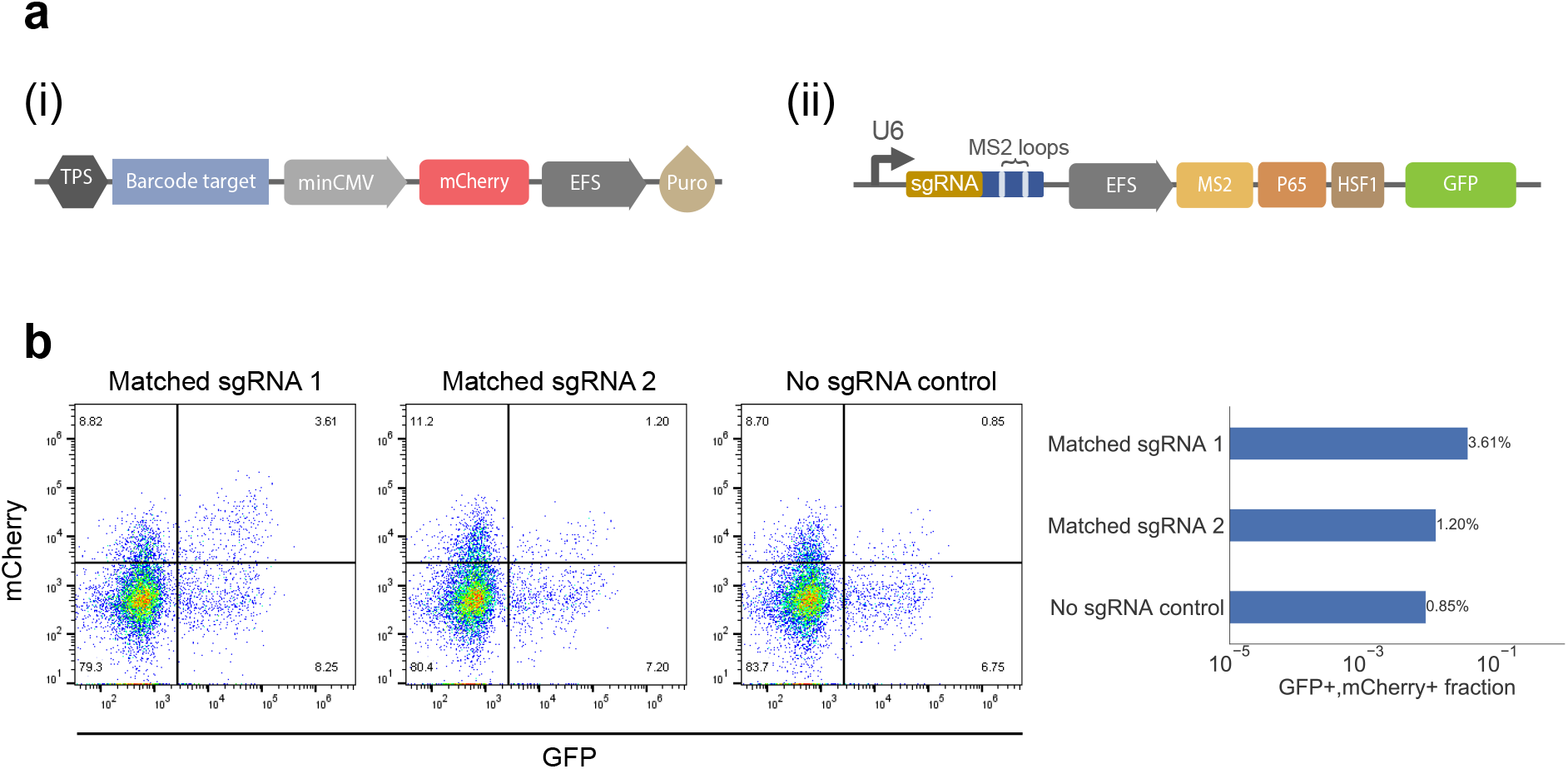
Transcriptional activation-based retrieval reporter. (a) (i) The reporter comprises transcriptional pause site (TPS), barcode target, minCMV promoter, mCherry fluorescent reporter and puro resistance marker. (ii) sgRNA with transcriptional activator. (a) The reporter selectively induces expression of mCherry in cells with matching sgRNA (Matched sgRNA 1 & Matched sgRNA 2), while cells without the sgRNA sequence exhibit a low level of mCherry expression (No sgRNA control).

**Fig. S3.**
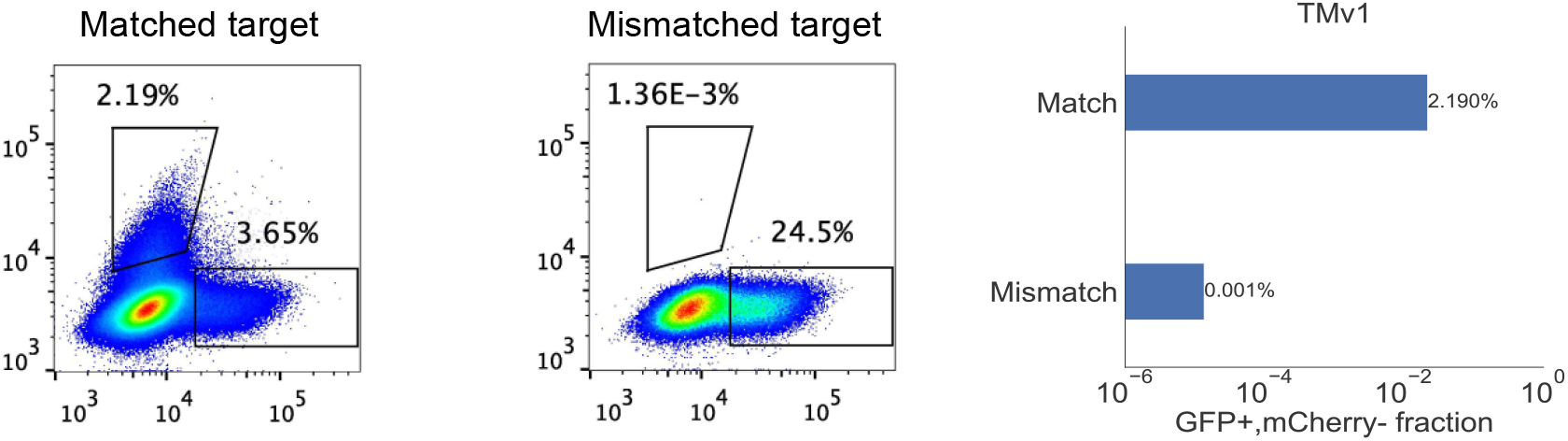
Specificity and sensitivity of the initial retrieval vector design (TMv1). TMv1 demonstrates high specificity, the background activation is around 1-4 in 100,000 cells (Matched sequence: GAGACCAGCAGAACCGACAA; Mismatched sequence: GCGCAACAGAGAGGGGAGCG).

**Fig. S4.**
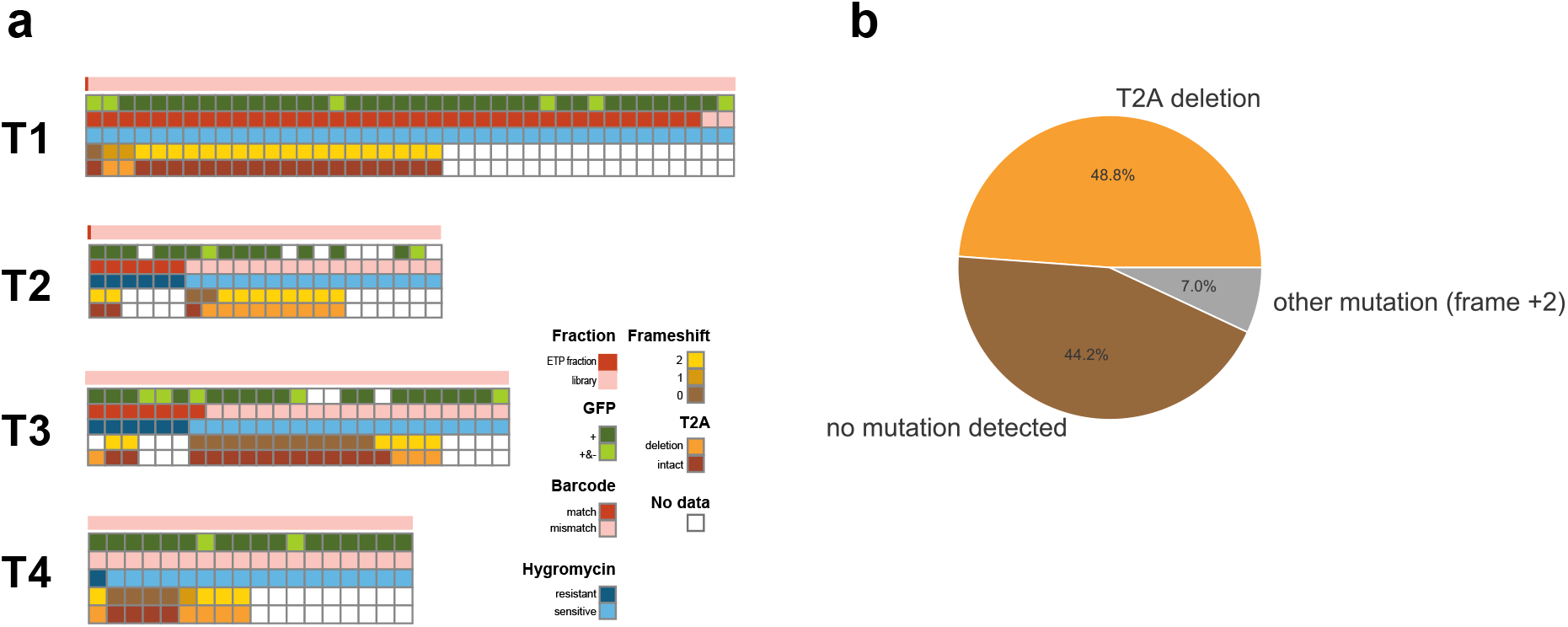
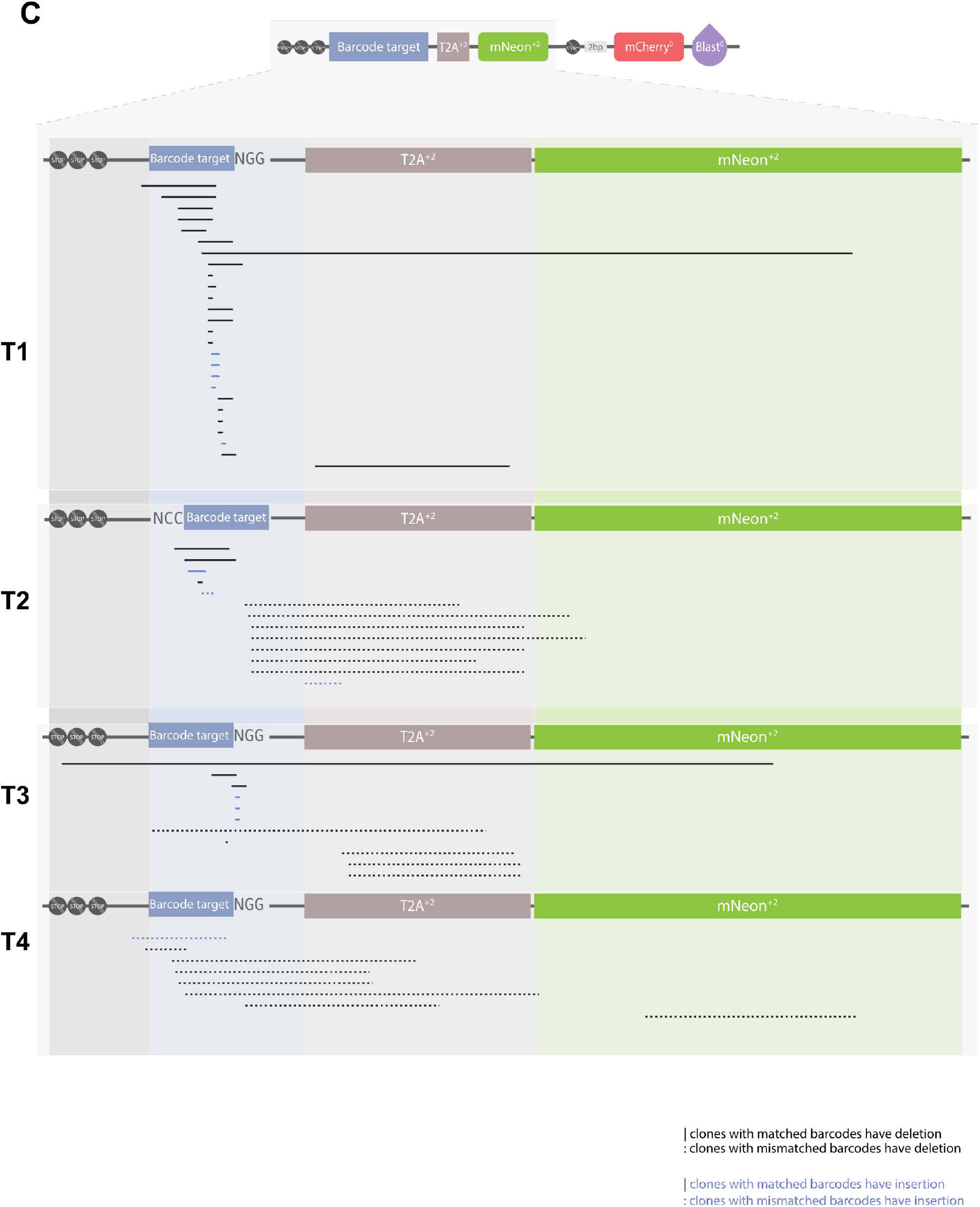
Characterization of retrieved clones. (a and b) Instances of retrieval of clones carrying incorrect barcodes are primarily explained not by FACS error, but by background mutations, in particular a stereotyped T2A linker deletion. (a) Sorted clones were analyzed by FACS for GFP expression (green), validated for barcode accuracy by NGS (red), validated for hygromycin sensitivity (blue) and Sanger sequenced to determine frameshift status (brown) and T2A (orange) deletion. (b) The types of background in the clones with mismatched barcodes are shown. T2A deletions account for 48.8% of background events. (c) Map of the retrieval vector, focusing on the targeting region. Each line represents a clone, with black and blue lines represent deletion and insertion regions, respectively and normal and dashed lines representing clones with matched and mismatched barcodes, respectively. The line depicts the location of deletion and the length is proportional to the size of deletion and insertion. Sanger sequencing data for each clone is provided as a SEQ file in Additional file 9: Sanger sequencing files.

**Fig S5.**
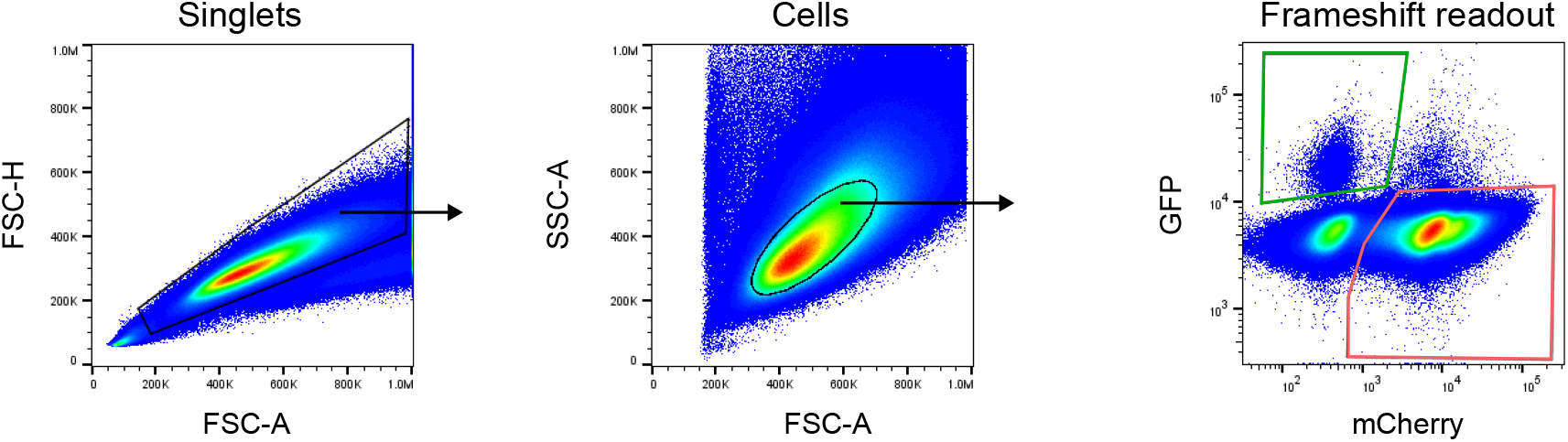
Gating strategy for analysis of cells with activated frameshift reporter. Representative flow plots for HeLa cells after frameshift mutation induced with Cas9.

**Table S1.**
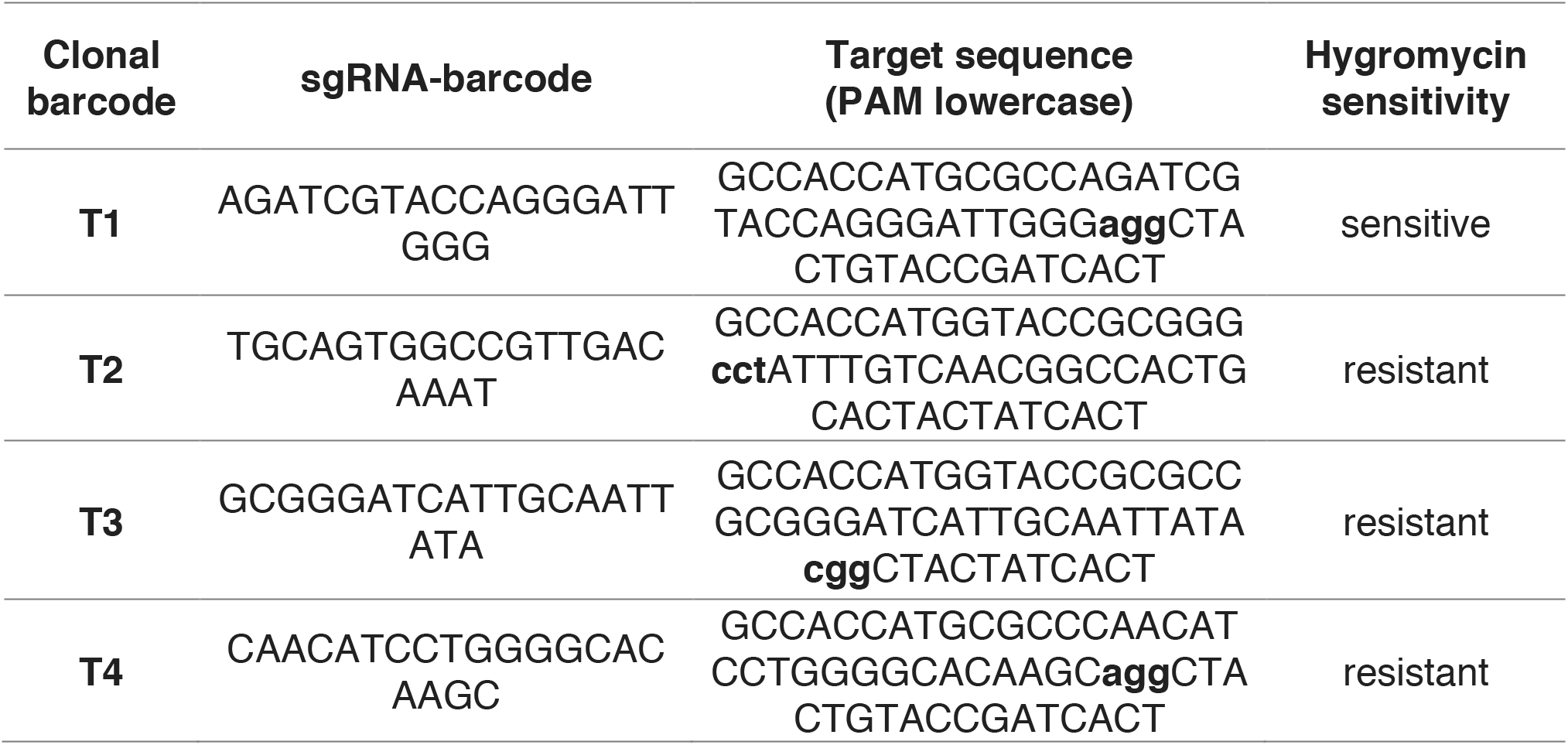
Table of clonal barcode candidates with their sgRNA-barcode sequences, corresponding target sequences and hygromycin sensitivity.

**Table S2.**
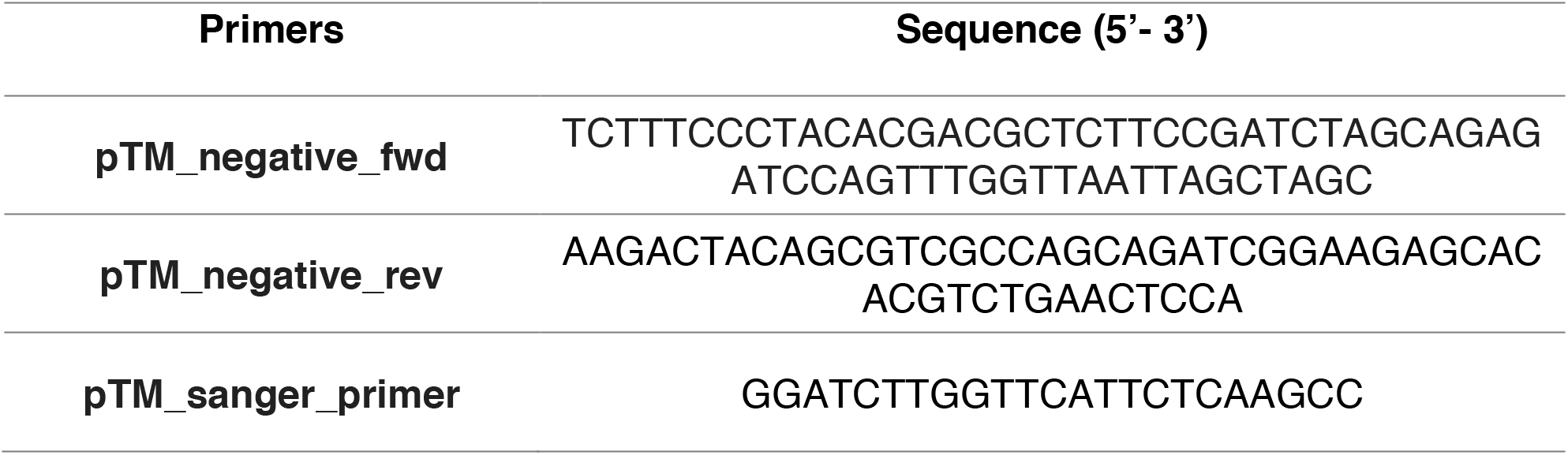
Table of primer sequences used for amplifying 2 kb-lentiviral transgene and for Sanger sequencing.

**Table S1. Table of sgRNA-barcode sequences from the D458 clonal tracking experiment.** The abundance of each sgRNA-barcode was calculated from normalized read counts and transformed by a base-10 logarithm (Method). Median rank and median values of each sgRNA-barcode in each condition across replicates are listed. Identified barcode sets including JQ1, DMSO and JQ1 & DMSO are listed in the last column.

**Table S2. Table of sgRNA-barcode sequences from the HeLa clonal tracking experiment.** The abundance of each sgRNA-barcode was calculated with normalized read counts and transformed by a base-10 logarithm (Method). Median ranks and median values of each sgRNA-barcode in each condition across replicates are listed. Identified barcode sets including HeLa and PBS are listed in the last column.

## Notes

https://drive.google.com/open?id=1cwTVK_0ttcihj7PprdkmDHNVfSGYOn8F

